# Acute oral toxicity and angiotensin-converting enzyme (ACE) inhibitory activity of the aqueous fractions of the methanol leaf extract of *Psychotria luzoniensis*

**DOI:** 10.1101/2025.10.14.681951

**Authors:** Ma. Danica Ines-Ramil, Kyoko Kobayashi, Kenroh Sasaki, Francisco M. Heralde

## Abstract

This study investigated the acute oral toxicity, *in vitro* angiotensin-converting enzyme (ACE) inhibitory activity, and metabolite profile of the aqueous fraction of the methanol leaf extract of *Psychotria luzoniensis* (PLAF), a plant endemic to the Philippines. ACE inhibition was evaluated in the crude extract, fractions, and sub-fractions through *IC*_50_ determination. The most bioactive fraction was further subjected to acute oral toxicity testing in mice at the limit dose of 2000 mg/kg body weight, while metabolite profiling was performed using UHPLC-QTOF-MS/MS. PLAF exhibited the strongest ACE inhibition (*IC*_50_ 34.38 ± 1.86 µg/ml), prompting subsequent fractionation. Among the sub-fractions (PLAF-1A - 1E), PLAF-1D demonstrated the most potent activity (*IC*_50_ 9.70 ± 0.10 µg/ml). Acute toxicity evaluation of PLAF revealed no mortality, behavioral abnormalities, or significant alterations in body weight, organ weights, serum biochemical parameters (SGPT, SGOT, BUN, creatinine, LDH), or histopathological changes in the liver, kidney, and heart. These findings indicate an *LD*_50_ greater than 2000 mg/kg. Metabolite profiling showed candidate flavonoids previously associated with ACE inhibition, including quercetin, kaempferol, rutin, and hyperin. Other putative compounds detected include 4-(3,4-dihydroxyphenyl)-7-hydroxy-2-oxo-2H-chromen-5-yl-β-D-glucopyranoside, juncein, and butin which are rarely reported in other *Psychotria* species. In this study, the ACE inhibitory activity and favorable toxicity profile of *P. luzoniensis* highlight its pharmacological potential to support further isolation of bioactive compounds and *in vivo* blood pressure studies.

## 1. INTRODUCTION

Hypertension remains a critical public health issue, particularly in developing countries like the Philippines, where its prevalence continues to rise. The World Health Organization (WHO) identifies hypertension as a leading cause of cardiovascular diseases, contributing significantly to global morbidity and mortality[1]. In the Philippines, low awareness, lack of treatment, and suboptimal blood pressure control exacerbates this issue, making the management of hypertension a pressing concern[2]. The severe complications of uncontrolled hypertension, such as heart disease, stroke, and kidney failure, emphasize the urgent need for more accessible and effective treatment options[3].

Blood pressure regulation is significantly influenced by the renin–angiotensin–aldosterone system (RAAS). The pathway involves angiotensin-converting enzyme (ACE), which first converts angiotensinogen into angiotensin I. Angiotensin I is then processed into angiotensin II, a molecule that not only constricts blood vessels to increase pressure but also triggers aldosterone release that leads to sodium and water retention and contributes to elevated blood pressure. Given the significant role of ACE in this system, inhibiting this enzyme is a primary strategy for managing blood pressure and reducing cardiovascular risk. However, synthetic ACE inhibitors, sometimes lead to kidney problems or elevated potassium levels [4],which can limit their use, especially in resource-limited settings.

*Psychotria luzoniensis*, (Cham. & Schltdl.) Fern.-Vill. from the family Rubiacea locally known as tagpong-gubat, is an endemic species with limited studies on its medicinal benefits. It has several synonyms, including *Coffea luzoniensis* Cham. & Schlecht., *Grumilea luzoniensis* (Cham. & Schltdl.) Merr., and *Paederia tacpo* Blanco [5]. The species is an erect, smooth shrub that grows 1.5–5 m in height, characterized by shiny, oblong-elliptic leaves, white terminal inflorescences, and fleshy, yellow to reddish fruits [5]. Although some species of *Psychotria* have been traditionally used in other countries for treating cardiovascular disorders due to their potential sympathetic blocking activity [6], there is limited phytochemical or pharmacological information available on the potential of *P. luzoniensis* for treating hypertension. *Psychotria* species are rich in alkaloids, coumarins, flavonoids, terpenoids, tannins, and cyclic peptides [7]. These compounds are known for their antihypertensive activity and have been used in traditional medicine to treat various disorders, including gastrointestinal, cardiovascular, and mental health conditions [6]. Given the chemotaxonomic and ethnomedicinal significance of the *Psychotria* genus, as well as its potential as a source of biologically active compounds, it is essential to investigate the antihypertensive activity of *P. luzoniensis*.

This research focuses on the leaf extracts of *P. luzoniensis* to explore its potential as a natural ACE inhibitor. Based on our previous report on the isolated compounds from the aqueous fraction of *Psychotria luzoniensis*, identified flavonoids such as quercetin glycosides and kaempferol derivatives, are known to exhibit antihypertensive activities [8]. Thus, this current study aims to further evaluate the therapeutic potential by assessing the acute toxicity and *in vitro* ACE inhibitory activity of its aqueous fraction. The investigation of *P. luzoniensis* is crucial, as limited studies have been conducted on this species, despite its potential as a source of biologically active compounds for direct use as drugs or as herbal remedies in the management of hypertension. This research aimed to contribute to support the global efforts to discover new, naturally derived solutions for managing hypertension, particularly in developing countries where access to conventional treatments is limited.

## 2. MATERIALS AND METHODS

### 2.1. Chemicals

ACE Kit—WST was purchased from Dojindo Corp. (Dojindo Laboratories, Kumamoto, Japan). ALT/GPT Activity Assay Kit (Colorimetric Method, Catalog No. E-BC-K235-S) and AST/SGOT Activity Assay Kit (Colorimetric Method, Catalog No. E-BC-K236-S) were from Elabscience^®^. All solvents (hexane and methanol) were of analytical grade. Other reagents were of research grade and were obtained from Wako Pure Chemical Industries, Ltd. (Osaka, Japan).

### 2.2. Ethical approval

All animal experiments were reviewed and approved by the Animal Experimental Committee of Tohoku Medical and Pharmaceutical University **(Approval No. A23073)**. The study was conducted in compliance with the university’s institutional ethical standards and international guidelines for the care and use of laboratory animals. Mice were euthanized following the recommendations outlined in the American Veterinary Medical Association (AVMA) Guidelines for the Euthanasia of Animals [9].

### 2.3. Plant materials

*Psychotria luzoniensis* was collected in Pasuquin, Ilocos Norte, Philippines, and identified at NUEBG (Northwestern University Ecological Park & Botanic Gardens). Voucher specimens were deposited and assigned as No. 14944 at the Herbarium of Northwestern Luzon (HNUL), Philippines. Mature leaf blades of each plant species were air-dried and homogenized for extraction and further analyses.

### 2.4. Extraction and liquid-liquid partitioning

One kilogram of powdered, air-dried leaves of *Psychotria luzoniensis* was macerated for 72 hours and extracted with methanol (MeOH, 500 ml × 6) at room temperature. The methanol extract was concentrated under reduced pressure using a rotary evaporator (Heidolph, Germany) at temperatures not exceeding 40°C and further concentrated using a speed vacuum (Eppendorf, Germany). Two-thirds of the crude extract was then suspended in water (1L) and partitioned with hexane (1L) to obtain aqueous and *n*-hexane-soluble fractions. The aqueous fraction was concentrated using a freeze dryer (Eyela, Tokyo, Japan), while the *n*-hexane fraction was concentrated sequentially with a rotary evaporator (Eyela, Tokyo, Japan) and a water bath concentrator (Biolab Scientific, Canada). Both fractions were screened for *in vitro* ACE I inhibitory activity. The bioactive aqueous fraction was further subjected to open-column chromatography over macroporous resin (DiaionTM HP20, Mitsubishi Chemical Corp., Tokyo, Japan), employing stepwise elution with solvent systems of increasing polarity (H_2_O: methanol, 10:0 → 8:2 → 6:4 → 2:8 → 0:10, v/v) followed by acetone. Complete solvent removal yielded six sub-fractions, designated as PLAF 1A to PLAF 1F, which were subsequently evaluated for *in vitro* ACE I inhibitory activity.

### 2.5. ACE inhibitory activity determination

The ACE inhibitory activities of the extract and sub-extracts were evaluated using the ACE Kit-WST (Dojindo Laboratories, Japan). The assay procedure was consistent with the assay kit technical manual provided by the manufacturer. Briefly, enzyme B was dissolved in 2 ml of deionized water, and 1.5 ml of this solution was then mixed with Enzyme A to prepare the enzyme sample. The indicator working solution was prepared by dissolving Enzyme C and the coenzyme separately in 3 ml of deionized water each, followed by combining 2.8 ml of the Enzyme C solution with 2.8 ml of the coenzyme solution.

The sample solutions were prepared by a five-fold serial dilution with deionized water, starting from an initial concentration of 300 µg/ml. For the assay setup, 20 µl of each sample solution was dispensed into the wells, along with two separate 20 µl aliquots of deionized water for blank 1 and blank 2. Substrate buffer (20 µl) was added to all wells, and an additional 20 µl of deionized water was placed in the 2 well. Subsequently, 20 µl of the enzyme working solution was added to each sample and blank 1 wells. The final concentration of the samples in the well were 100 µg/ml to 0.16 µg/ml. To correct for background absorbance due to the inherent color of the extract, a separate sample blank was prepared for each concentration, consisting of 20 µl of the sample solution diluted with 240 µl deionized water. The microplate was incubated at 37°C for 1 hour. After incubation, 200 µl of the indicator working solution was added to each well except for the sample blank wells, and the plate was further incubated at room temperature for 10 minutes. Absorbance was measured at 450 nm using a microplate reader (Corona Absorption Grating Microplate Reader SH-1300Lab, Corona Electric Co., Ltd., Ibaraki, Japan). The percentage of ACE inhibitory activity was calculated using the following equation:

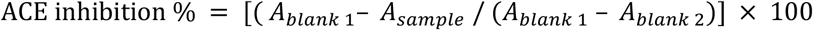

where *A*_*blank* 1_ represented positive control (without ACE inhibition); *A*_*blank* 2_ was the absorbance of the reagent blank, while *A*_*sample*_ was the absorbance of the sample. The concentration inducing 50% inhibition or the *IC*_50_ of the samples, was also determined using non-linear regression. All assays were carried out in three independent experiments, each performed with three replicates per sample at different concentrations. For each experiment, *IC*_50_ values were calculated from the replicates, and the mean *IC*_50_ was obtained. The overall *IC*_50_ value was then determined as the average of the means from the three independent experiments, and results are expressed as mean ± SD. Captopril was used as standard.

### 2.6. UHPLC-QTOF-MS/MS metabolite profiling

Metabolite profiling was performed on a UHPLC system (Nexera X2, Shimadzu, Kyoto, Japan) coupled to a quadrupole time-of-flight mass spectrometer (LCMS-9050, Shimadzu, Canby, OR, USA).

Chromatographic separation was achieved using a Waters HSS T3 column (2.1 mm ID × 100 mm, 1.8 µm particle size). The mobile phases consisted of 0.1% formic acid in water (solvent A) and 0.1% formic acid in acetonitrile (solvent B).

The mass spectrometer was operated in electrospray ionization (ESI) mode with polarity switching between positive and negative ion modes. Mass spectrometric data were acquired using a data-dependent acquisition (DDA) method over a run time of 0.00–12.00 min. Each acquisition cycle included one full MS scan (m/z 80–1000) followed by 20 MS/MS events, each targeting a different precursor ion with a product ion scan range of m/z 40–1000. The collision energy (CE) was set to 30 V with a spread of ±25 V, and each MS/MS event lasted 0.040 s. The Q1 resolution was set to *Unit*, and MS/MS triggering required a high-intensity threshold. The interface voltage was maintained at 1.50 kV, while voltages for the DL, Qarray, and Q1 Pre were automatically sourced from the instrument tuning file.

Sample preparation was carried out by weighing 1 mg of each sample, dissolving it in 1 mL of methanol, followed by sonication for 10 min and filtration through a 0.22 µm PVDF syringe filter prior to injection.

Data processing was performed using Insight Discovery software, which matched acquired spectra against a user-provided screening list, the NIST 2023 spectral library, and an in-house metabolite library. Additional data mining was performed with Insight Explore software for formula prediction (accurate mass, isotope patterns, adduct types, elemental constraints). Compound identification and confirmation were based on the following criteria: exact mass tolerance (±3 ppm), retention time (±0.5 min), and MS/MS library match score index (>80) including isotope and product ion pattern comparison. Structural validation of detected compounds was performed using the MS/MS Assign module, with cross-referencing against public databases such as PubChem and ChemSpider.

### 2.7. Acute oral toxicity study

All experiments involving test animals were approved by the Animal Experimental Committee of Tohoku Medical and Pharmaceutical University (Approval No. A23073), and the experimental procedures were conducted in accordance with the ethical guidelines of the university.

Twelve female ICR mice (8 weeks old) purchased from Japan SLS Inc. were housed and maintained under standardized conditions of temperature (25±1°C) and humidity (55±5%) in a light cycle room. The mice were allowed to acclimatize for a week and fed with standard chow (CE-2; CLEA Japan, Inc., Tokyo, Japan).

The acute oral toxicity study was performed according to OECD Guideline 423, using female mice, as recommended, to minimize interspecies variation and reduce the number of animals used. Briefly, 12 ICR mice were randomly divided into two groups (control and PLAF *n*=6 in each group). First three mice from each group were fasted for 12h before the experiment with free access to water until 3-4 hours before the experiment. Each mouse (I–III) in both the PLAF-treated and control groups was orally administered either a limit dose of 2000 mg/kg body weight of the extract or distilled water at a volume equivalent to 1% of body weight (0.1 mL per 10 g), respectively. The mice were observed for toxic symptoms within the first four hours after administration, food was withheld only 1-2 hours during that time. No mortality was recorded among the initial three mice in each group within 48 hours of treatment. Consequently, an additional three mice were administered the same dose of 2000 mg/kg body weight. As no deaths occurred in either set of animals, no further testing was required. Behavioral changes, toxic effects, and mortality were observed and recorded daily in all groups for 14 days after dosing. Body weights of mice in each group were recorded before oral administration and at least three times during the administration period (including at the end of the study) to calculate the weight alterations.

### 2.8. Measurements of organ weight, macroscopic observation and histopathological examination

Mice were euthanized under isoflurane (FUJIFILM Wako Pure Chemical Corporation, Japan) anesthesia at the end of experiment. Blood samples were collected via cardiac puncture. Blood samples were allowed to clot at room temperature for 30 minutes and were then centrifuged at 3000 × g for 10 minutes at 4 °C to obtain serum. The resulting serum was carefully aliquoted and stored at –80 °C until further biochemical analysis. Vital organs, including the heart, the liver, and the kidney, were removed, washed with iced-cold phosphate-buffered saline (FUJIFILM, Wako Pure Chemical Corporation, Osaka, Japan), dried with laboratory lint free wipes and weighed using analytical balance. The relative weight of each organ was be calculated using the following formula:

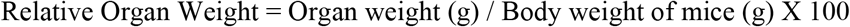

The morphology was macroscopically observed for signs of toxicity. Samples of organs isolated from each mouse were fixed in 10% formalin neutral buffer solution (FUJIFILM, Wako Pure Chemical Corporation, Osaka, Japan), dehydrated by serial ethanol solution and enclosed with paraffin wax. Micrometer sections (5 μm) cut with microtome were stained with hematoxylin and eosin and were examined under a light microscope, photomicrographs of the samples were recorded.

### 2.9. Biochemical analyses

#### 2.9.1. Liver Function Markers: ALT and AST

Serum alanine aminotransferase (ALT/SGPT) and aspartate aminotransferase (AST/SGOT) activities were measured using commercially available colorimetric assay kits (Elabscience; ALT, Cat. No. E-BC-K235-S; AST, Cat. No. E-BC-K236-S). Assays were adapted for 96-well microplate format following the manufacturer’s instructions. Briefly, 25 µl of serum sample, blank, or pyruvate standard was mixed with 50 µl of substrate working solution and incubated at 37°C for 30 minutes. After incubation, 25 µl of chromogenic reagent (2,4-dinitrophenylhydrazine, DNPH) was added, followed by an additional 20-minute incubation. Reactions were terminated with 125 µl of alkali reagent, and absorbance was measured at 510 nm using a microplate reader. Enzyme activities were calculated based on standard curves generated from serial dilutions of a 2 mmol/L sodium pyruvate standard and expressed as units per liter (U/L). All measurements were performed in duplicate.

#### 2.9.2. Renal Function Markers: Creatinine and BUN

Serum creatinine and blood urea nitrogen (BUN) levels were determined using the Roche Cobas C 311 Clinical Chemistry Analyzer (Roche Diagnostics, Germany), which employs validated enzymatic and kinetic methods. Creatinine concentration was measured using an enzymatic colorimetric assay (CREP2 method), with absorbance read at 546 nm. BUN was quantified using a kinetic UV assay (UREA2 method) based on the urease–GLDH reaction, with monitoring of NADH absorbance decrease at 340 nm. Results were automatically calculated and expressed in milligrams per deciliter (mg/dL).

#### 2.9.3. Tissue Injury Marker: LDH

Lactate dehydrogenase (LDH) activity was measured using the Roche Cobas C 311 Analyzer with the LDHI2 assay. LDH activity was assessed based on the enzymatic conversion of L-lactate to pyruvate, coupled with the reduction of NAD^+^ to NADH, with absorbance continuously monitored at 340 nm. LDH activity was automatically calculated from the rate of absorbance change and reported in international units per liter (IU/L).

##### Statistical Treatment of Data

Normality of data was verified using the Shapiro–Wilk test. Values are presented as mean ± SD with 95% confidence intervals, and significance was set at *p* < 0.05. Group differences were assessed by one-way ANOVA, with Tukey’s post hoc test for equal variances or Welch’s ANOVA followed by Dunnett’s T3 for unequal variances (Brown–Forsythe test, *p* < 0.05). Comparisons between two groups were performed using the two-tailed unpaired t-test. Statistical analyses were conducted using GraphPad Prism® software version 10.6.1 (GraphPad Software, Boston, Massachusetts USA).

## 3. RESULTS

### 3.1. ACE-I inhibitory activity of the methanol leaf extract and subsequent fractions of *P. luzoniensis*

#### 3.1.1. In vitro ACE inhibitory activity of PLC, PLAF, and PLHF

*In vitro* angiotensin-converting enzyme (ACE 1) inhibitory assay was conducted on polarity-based fractions of the methanol crude extract. This bioassay-guided approach enabled the comparison of polar and non-polar fractions in terms of ACE inhibition, serving as a predictive measure of their pharmacological relevance. The fraction demonstrating the highest inhibitory activity was subjected to further fractionation and an acute oral toxicity evaluation.

*Psychotria luzoniensis* methanol crude extract (PLC) and aqueous fraction (PLAF) showed similar ACE inhibitory activities (adjusted *p-*value = 0.9984), with *IC*_50_ values of 40.75 ± 0.25 µg/ml and 34.38 ± 1.86 µg/ml, respectively. Both PLC and PLAF exhibited significantly more potent ACE inhibition compared to the *n*-hexane fraction (PLHF), which had an *IC*_50_value of 2,786.75 ± 1,036.33 µg/ml (*p* < 0.0001). These results indicated that ACE inhibitory compounds from *Psychotria luzoniensis* are predominantly polar. Captopril, used as the reference standard, evaluated at concentrations of 2.5–40 ng/mL obtained an *IC*_50_ of 12.5 ng/ml.

Based on the *in vitro* ACE inhibitory activity and its ethnopharmacological relevance, the PLAF was subjected for further fractionation and *in vivo* acute toxicity studies. The plant, being endemic and limited in availability, was evaluated at the fraction and sub-fraction levels to conserve material while enabling pharmacological investigation. Furthermore, its traditional use as a decoction justified the investigation of the aqueous extract in its near-native form.

#### 3.1.2. ACE-I inhibitory activity of bioassay-guided sub-fractions of PLAF

To identify the most bioactive fraction from the leaf extract of *Psychotria luzoniensis*, we evaluated the ACE inhibitory activity of six fractions (1A–1F) across four concentrations (0.8, 4, 20, and 100 µg/ml). As shown in Figure 2, most fractions exhibited a concentration-dependent inhibition profile, although 1A, 1B, and 1F showed limited variation across doses. Two-way ANOVA confirmed significant effects of both concentration and fraction on ACE inhibition (*p* < 0.0001), with a significant interaction between the two factors. PLAF-1D consistently demonstrated the highest ACE inhibition, with mean values of 97.95 ± 0.08%, 70.39 ± 0.20%, 20.54 ± 0.41%, and 19.31 ± 0.56% at 100, 20, 4, and 0.8 µg/ml, respectively. Tukey’s multiple comparisons test indicated that PLAF-1D was significantly more active than most other fractions at each concentration tested (*p* < 0.05).

**Figure 1.**
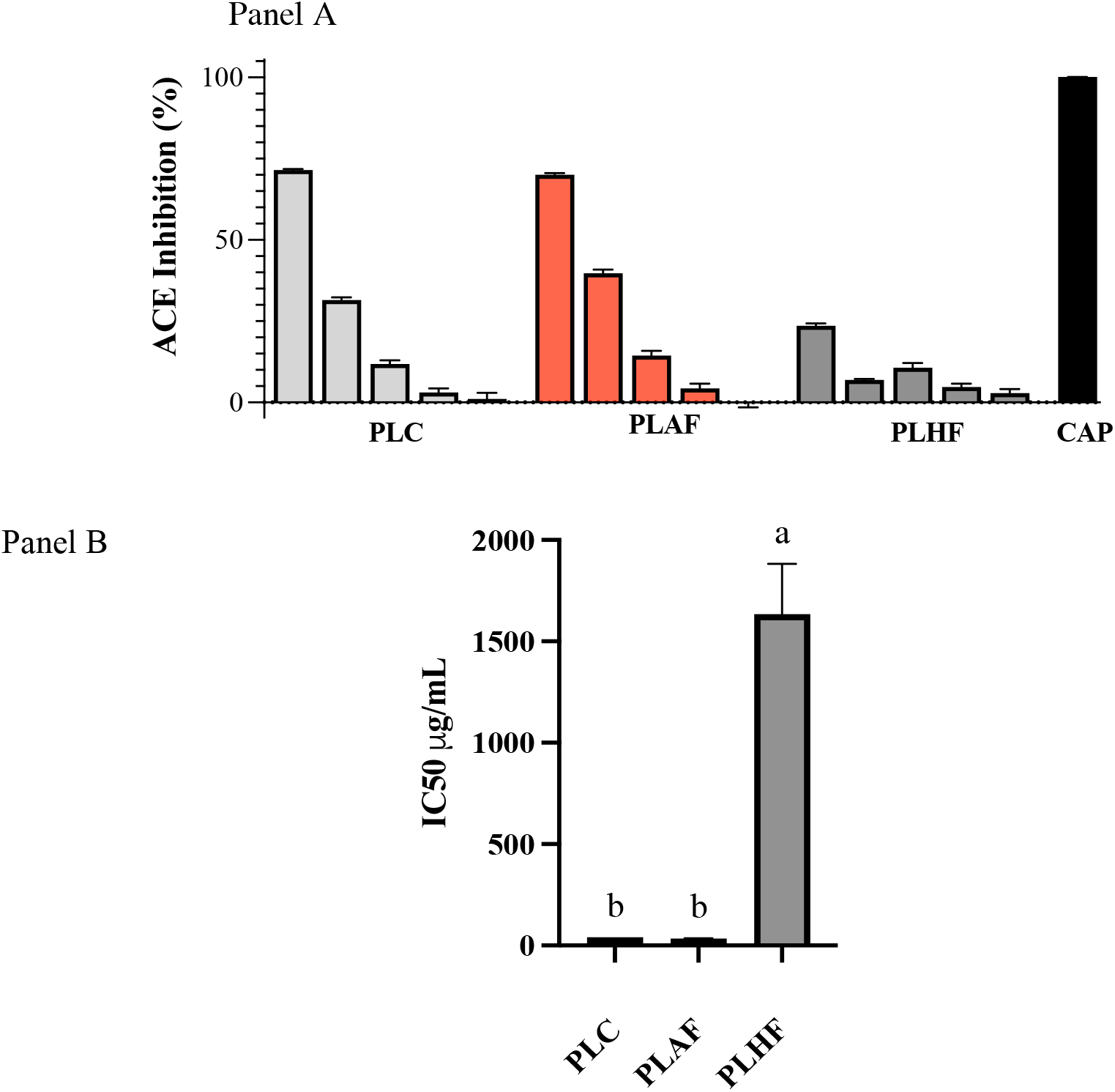
Concentration-Dependent ACE Inhibition of *Psychotria luzoniensis* Extract and Fractions (A) and Corresponding IC_50_ Values (B) **Note: (Panel A)** In vitro angiotensin-converting enzyme (ACE) inhibitory activity (%) of Psychotria luzoniensis samples following concentration dependency at 100, 20, 4, and 0.8 µg/ml. **(Panel B)** The bars represent the mean IC_50_ values of the crude extract and fractions (µg/mL). Data are expressed as mean ± SD from three independent experiments (n = 3), each performed in triplicate. CAP exhibited an IC_50_ of 12.5 ng/ml. Statistical analysis was performed using ordinary one-way ANOVA followed by Tukey’s post hoc test. All pairwise comparisons between extract groups were statistically significant at p < 0.0001.

**Figure 2.**
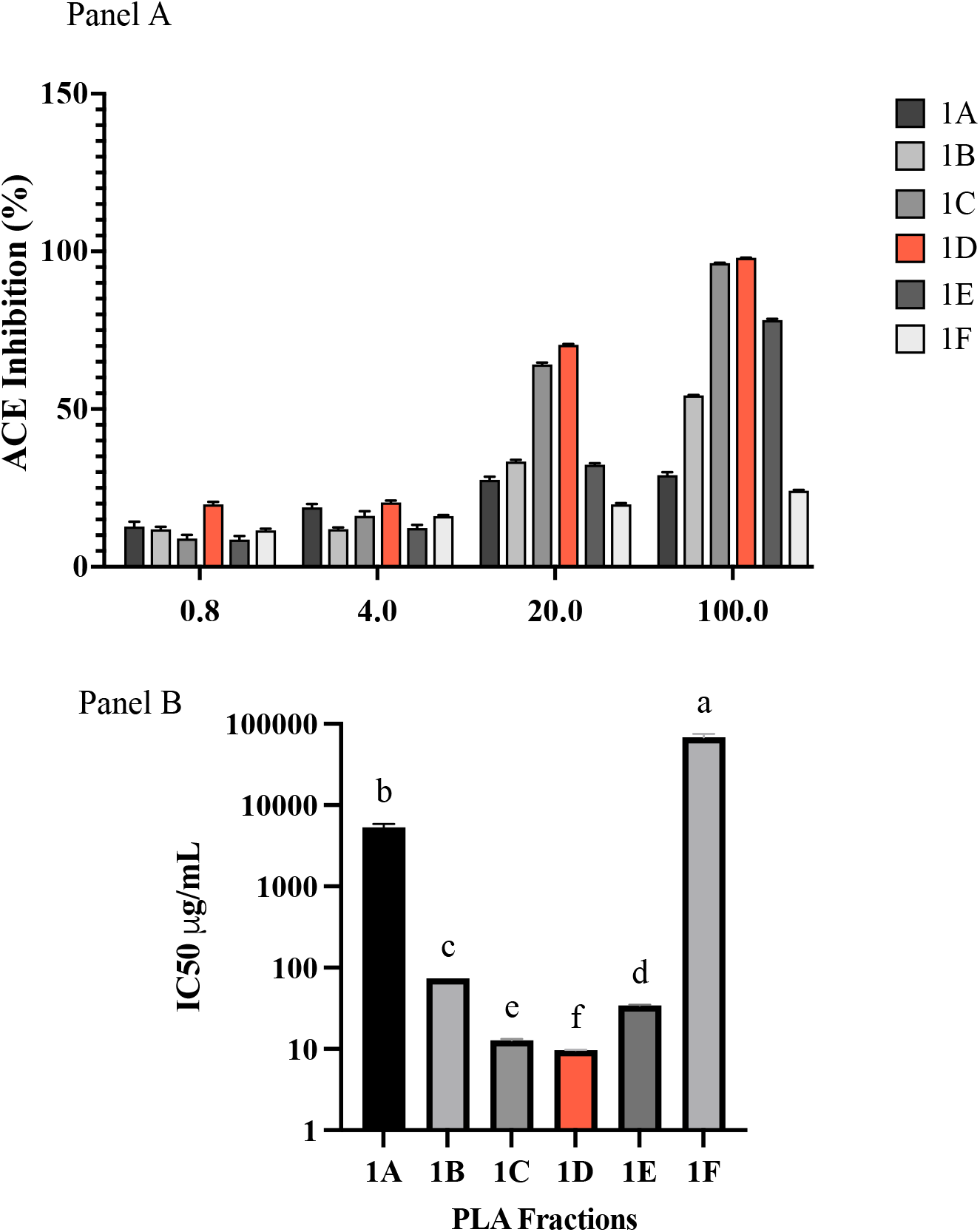
Concentration-Dependent ACE Inhibition and IC_50_ Values of the Aqueous Fractions of *Psychotria luzoniensis*. **Note: (Panel A)** Concentration-dependent and fraction-dependent ACE inhibitory activity (%) of six fractions of Psychotria luzoniensis at 0.8, 4, 20, and 100 µg/ml confirmed by two-way ANOVA (p < 0.0001) followed by Tukey’s multiple comparison. (**Panel B)**. Bars represent the **c**orresponding mean IC_50_ values (µg/ml) of the fractions calculated by non-linear regression using a four-parameter logistic model. Data are expressed as mean ± SD from three independent experiments (n = 3), each performed in triplicate. CAP was used as standard exhibited an IC_50_ of 12.5 ng/ml. Statistical analysis was conducted using Welch’s ANOVA due to unequal variances (Brown-Forsythe test, p < 0.05), followed by Dunnett’s T3 multiple comparisons test.

To compare the potency of the fractions, *IC*_50_ values were determined through nonlinear regression. Welch’s ANOVA revealed significant differences among the fractions (*p* < 0.0001), and Dunnett’s T3 test identified fraction PLAF-1D as having the lowest *IC*_50_ (9.70 ± 0.10 µg/ml), followed by 1C (12.80 ± 0.41 µg/ml) and 1E (34.32 ± 0.58 µg/ml). Fractions 1A and 1F exhibited substantially higher *IC*_50_ values at 5332.89 ± 532.41 µg/ml and 68666.26 ± 7050.86 µg/ml, respectively, indicating relatively weak ACE inhibitory effects. These findings highlight fraction PLAF-1D as the most active sample based on both inhibition percentages and *IC*_50_ values. Captopril showed an *IC*_50_ of 12.5 ng/ml.

### 3.2. UHPLC-QTOF-MS/MS metabolite profile

LC-MS profiling of the bioactive PLAF revealed six putatively annotated compounds, including flavonol glycosides, free flavonoids, and other polar secondary metabolites. Major glycosylated flavonoids identified were rutin and hyperin, exhibiting fragmentation indicative of sugar moieties and their aglycone cores. Free quercetin was also detected, characterized by their key fragments. Additionally, choline, quercetin-3-O-β-D-xylopyranosyl-(1→6)-β-D-glucopyranoside, and 4′,5,7-trihydroxyflavone-β-D-glucopyranoside were identified, each supported by accurate mass measurements (mass error ±3 ppm).

Also, UHPLC-QTOF-MS/MS analysis of the most active sub-fraction (PLAF-1D) from *Psychotria luzoniensis* identified nine putatively annotated compounds, primarily flavonoids and phenolic glycosides. Primary metabolites detected were hyperin, quercetin, rutin, kaempferol, and kaempferol 3-β-D-galactoside. Other compounds included cyanidin 3-O-lathyroside cation, juncein, butin, and loliolide. The identifications were supported by their characteristic fragmentation patterns and accurate mass measurements with low mass errors (–0.601 to –1.167 ppm).

### 3.3. Acute oral toxicity

PLAF was assessed for acute oral toxicity in female ICR mice following OECD Guideline 423 as recommended to minimize interspecies variation and reduce the number of animals used. A single oral dose of 2000 mg/kg body weight was administered, and observations were conducted over 14 days. Parameters evaluated included determination of *LD*_50_, behavioral and signs of toxicity observations, gross necropsy of vital organs (liver, heart, and kidney), organ weights, microscopic examination, and biochemical assays for serum glutamic-pyruvic transaminase (SGPT), serum glutamic-oxaloacetic transaminase (SGOT), blood urea nitrogen (BUN), creatinine, and lactate dehydrogenase (LDH) levels to detect potential liver, kidney, and heart damage. The following results provide essential safety data on *Psychotria luzoniensis*, supporting and guiding further pharmacological and toxicological evaluations.

#### 3.3.1. Behavioral pattern, mortality and body weight

Following oral administration of PLAF at 2000 mg/kg BW, animals were closely monitored for three hours and regularly observed daily until day 14. No significant signs of acute toxicity or mortality were recorded as summarized in Table 1. Mild drowsiness and calming behavior were noted within the first 30 minutes, with animals subsequently falling asleep. These transient effects resolved without any abnormalities in external appearance.

As shown in Figure 3, mean body weights (BW) of mice treated with *Psychotria luzoniensis* aqueous extract (PLAF, 2000 mg/kg BW) remained comparable to controls over 14 days (*n* = 6, mean ± SD). No significant differences were observed at baseline and Day 14 (*p* > 0.05), although a transient BW reduction occurred on Day 10 in the PLAF group (*p* = 0.0012). Control mice showed significant BW gains between Days 0–3 and 7–10 (*p* < 0.05) and a moderate overall increase from Day 0 to Day 14 (*p* = 0.0214). PLAF-treated mice showed no significant changes until Day 10 but had a moderate BW increase by Day 14 (*p* = 0.0160). These results indicate that PLAF at 2000 mg/kg BW did not induce sustained weight loss or toxicity, with the temporary decrease likely reflecting a minor, reversible physiological effect.

**Figure 3.**
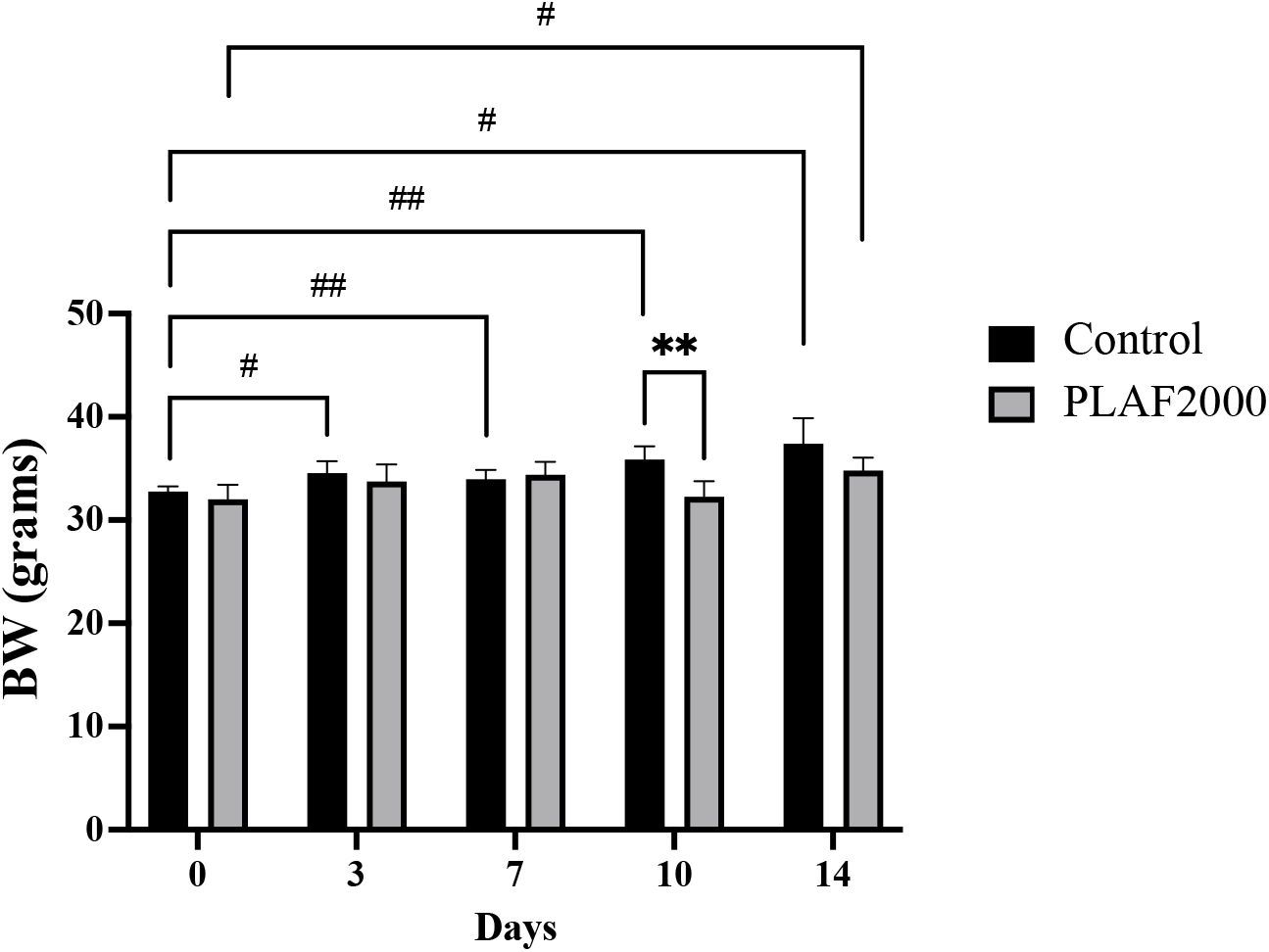
Changes in body weight of mice after a single oral dose of *Psychotria luzoniensis* aqueous extract (PLAF, 2000 mg/kg BW). **Notes:** Body weights were measured on Days 0, 3, 7, 10, and 14. Data are expressed as mean ± SD (n = 6 per group). *p < 0.05 vs. control; #p < 0.05 vs. Days within the same group. Statistical analysis was conducted using two-way ANOVA with Tukey’s post hoc test.

#### 3.3.2. Relative organ weights

The macroscopic examination of major organs, including the liver, kidneys, and heart, revealed no observable pathological lesions in either the treated or control groups. Gross morphological assessment revealed normal organ architecture in all experimental mice, with no detectable deviations from the control, indicating the absence of treatment-related toxicity.

The relative organ weights of vital organs (liver, heart, and kidneys) from mice treated with the aqueous extract of *Psychotria luzoniensis* at 2000 mg/kg showed no significant differences compared to the control group (p > 0.05) as shown in Figure 4. This suggests that acute administration of the extract did not cause noticeable organ enlargement, atrophy, or pathological changes within the experimental period.

**Figure 4.**
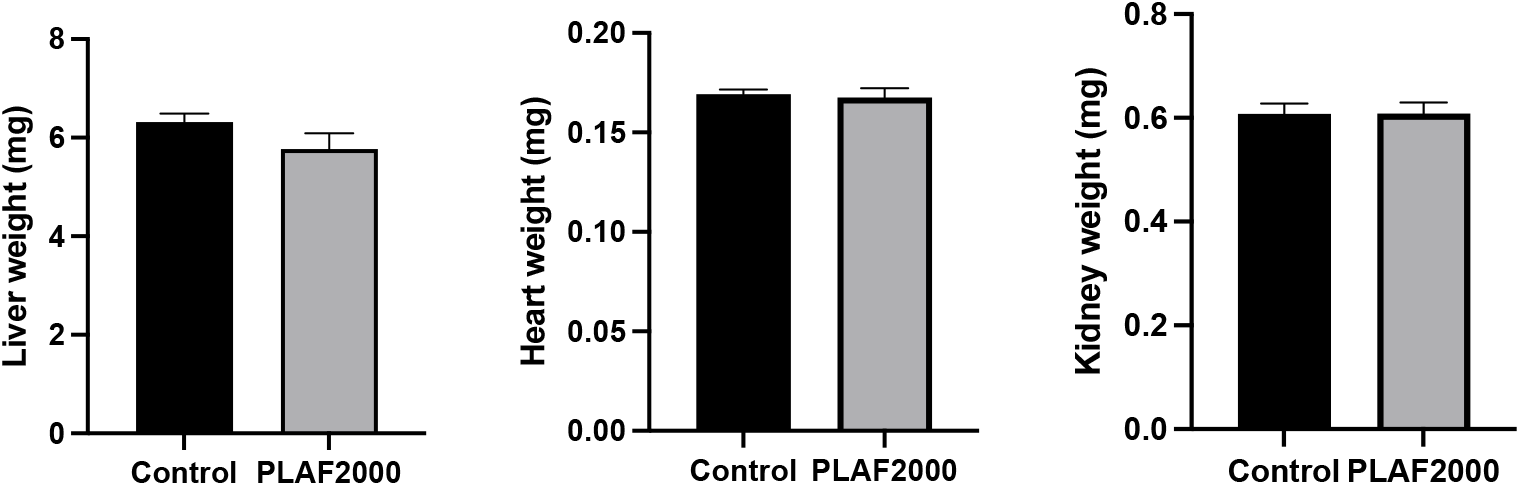
Effect of *Psychotria luzoniensis* aqueous extract (2000 mg/kg BW) on relative organ weights in mice. **Notes:** Relative weights of the liver, heart, and kidneys were measured in mice administered a single oral dose of PLAF (2000 mg/kg BW) compared to untreated controls. No significant differences were observed between groups for any organ (p > 0.05). Data are presented as mean ± SD (n = 6); unpaired t-test.

#### 3.3.3. Histology of liver, heart, and kidney

Although macroscopic and organ weight evaluations revealed no significant differences between control and PLAF2000-treated groups, the observed significant reduction in body weight of animals treated with PLAF at 2000 mg/kg BW on day 10 supported further histopathological assessment to identify potential microscopic tissue changes or subclinical toxic effects.

As shown in Figure 5, microscopic observation indicated no noticeable differences between control and PLAF-treated groups. The liver, heart, and kidney tissues from extract-treated mice exhibited normal cellular architecture with no adverse alterations under light microscopy. The cell structures evaluated included liver components (central vein, sinusoids, hepatocytes), cardiac tissues (muscle fibers, connective tissue), and kidney structures (glomerulus, podocytes, Bowman’s capsule, capillaries, proximal convoluted tubules, tubular lumens).

**Figure 5.**
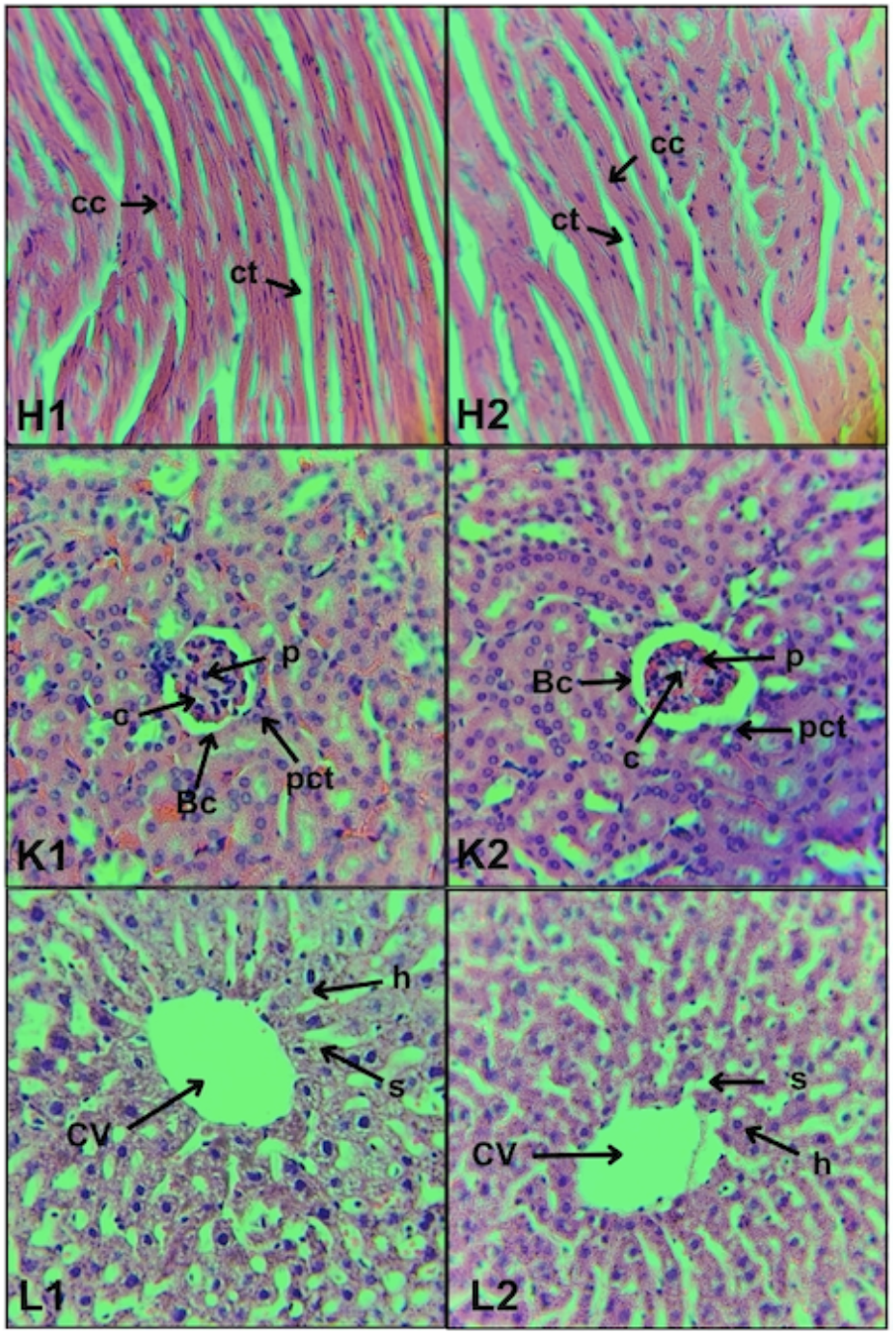
Representative photomicrographs of histological preparation of heart, kidney and liver controls (1) and treated (2) with aqueous extract of *Psychotria luzonienis*. **Notes:** Representative photomicrographs of histological preparation (0.5 µm), stained with hematoxylin and eosin of liver (L1, L2) heart (H1, H2), and kidney (K1, K2) of rats controls (1) and treated (2) with aqueous extract of Psychotria luzoniensis under a compound microscope at a 40x magnification. Abbreviation: cc (cardiac muscle cell), ct (connective tissue), g (glomerulus), p (podocyte), Bc (Bowman’s capsule), c (capillary), pct (proximal convoluted tubule), cv (central vein), s (sinusoids), h (hepatocyte).

#### 3.3.4. Biochemical analysis

Serum biochemical markers are critical indicators of systemic toxicity, particularly in relation to hepatic and renal function, as well as general tissue integrity. The biochemical analysis revealed no statistically significant differences (p > 0.05) in serum levels of SGPT, SGOT, BUN, Creatinine, and LDH between the control and PLAF2000 mg/kg BW-treated groups, as shown in Figure 6. The measured parameters remained within normal physiological ranges, indicating that the acute administration of the extract at 2000 mg/kg did not result in hepatic or renal impairment or general tissue damage.

**Figure 6.**
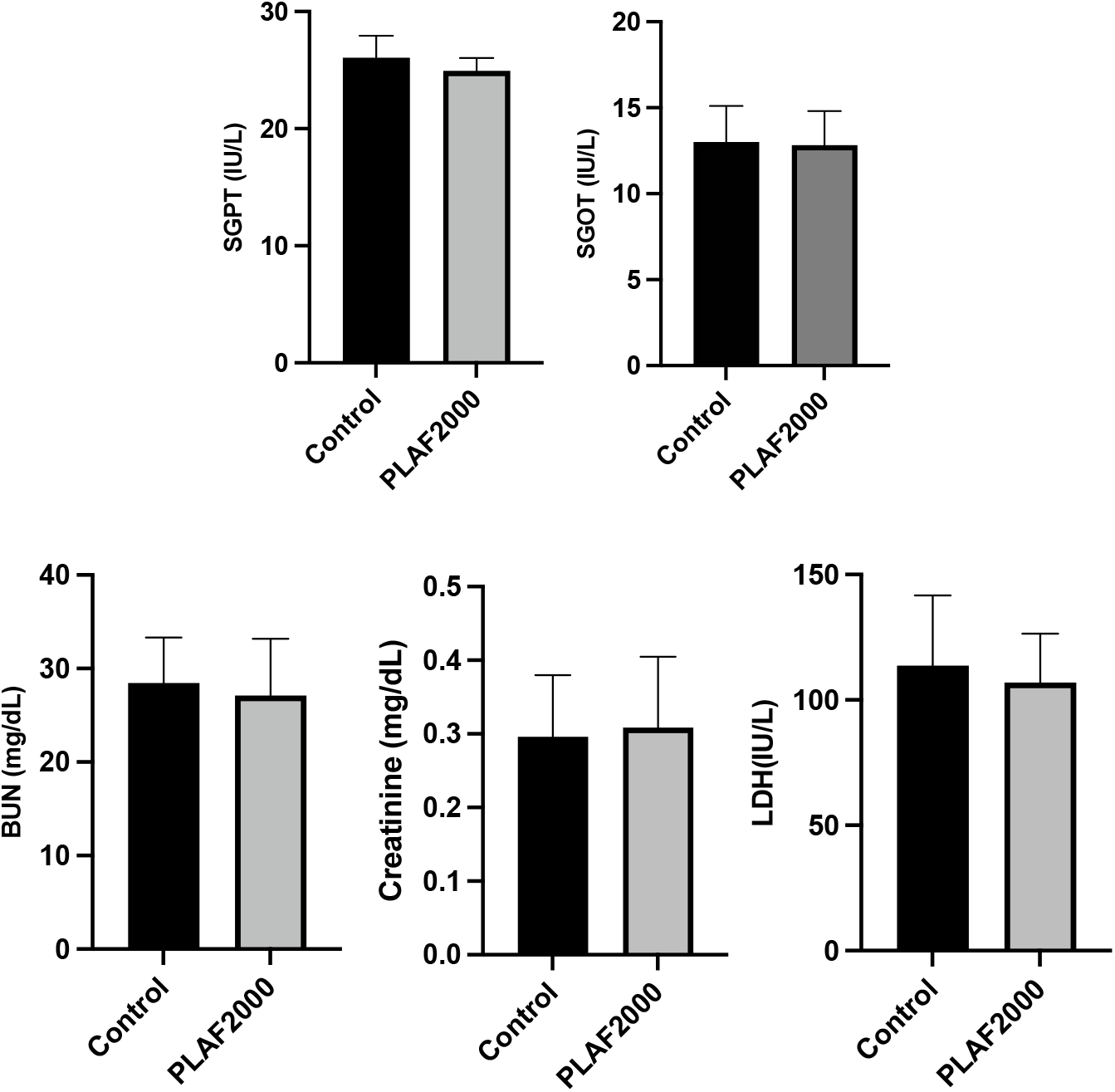
Serum biochemical parameters in mice treated with *Psychotria luzoniensis* aqueous extract (2000 mg/kg BW). **Note:** No significant differences (p > 0.05) were observed in SGPT, SGOT, BUN, creatinine, and LDH levels between the PLAF-treated and control groups. Data are presented as mean ± SD (n = 6); unpaired t-test.

The absence of significant alterations in SGPT and SGOT levels indicates that PLAF2000 administration did not elicit liver damage. The similar BUN and creatinine levels between the treated and control groups also confirm that renal function remained unaffected. Normal LDH levels indicate that there was no detectable generalized tissue or organ injury.

## 4. DISCUSSION

*Psychotria luzoniensis*, an endemic and poorly studied Philippine plant, was investigated in this study as a potential source of natural antihypertensive agents. It was selected based on its chemotaxonomic similarity to other *Psychotria* species with known cardiovascular activity. Activity-guided fractionation, followed by chemical profiling to identify putative ACE I inhibitory compounds, and acute toxicity testing facilitated assessment of the plant’s pharmacological potential and safety profile.

ACE I inhibition is a key pharmacological mechanism in hypertension management, as it suppresses the formation of angiotensin II, to promote vasodilation and reducing blood pressure. *In vitro* ACE 1 inhibitory activity showed that between aqueous (PLAF) and *n*-hexane (PLHF) fractions —PLAF exhibited higher inhibitory activity, indicated by the lowest *IC*_50_value of 34.39 ± 0.88 µg/ml with no significant difference with the parent extract PLC (*IC*_50_ = 40.69 ± 1.55 µg/ml), and was selected for further sub-fractionation. Fractionation of the aqueous portion using macroporous resin chromatography identified PLAF-1D, eluted with 80% methanol, as the most active fraction. This indicates that the ACE I inhibitory activity is associated with polar to moderately polar constituents concentrated in the methanolic fraction.

LC-MS profiling of PLAF and its sub-fraction PLAF-1D, revealed both overlapping and distinct metabolite profiles. Rutin, hyperin, and quercetin were consistently detected in both. Metabolites such as choline, quercetin-3-O-β-D-xylopyranosyl-(1→6)-β-D-glucopyranoside, and 4′,5,7-trihydroxyflavone-β-D-glucopyranoside were identified only in PLAF and were no longer detected in PLAF-1D, possibly due to their distribution across other sub-fractions during chromatographic separation. In contrast, PLAF-1D exhibited additional metabolites, including kaempferol, kaempferol 3-β-D-galactoside, cyanidin 3-O-lathyroside cation, juncein, butin, and loliolide. The higher number of compounds detected in PLAF-1D can be attributed to the concentration of specific constituents through sub-fractionation and enhanced resolution by UHPLC, which allowed more effective separation of co-eluting analytes present in the parent fraction.

Some putatively identified compounds have limited studies supporting their ACE inhibitory activity. However, kaempferol, quercetin, rutin, and hyperin have been reported to modulate angiotensin-converting enzyme (ACE), a key regulator of blood pressure. Kaempferol inhibits ACE through Zn^2+^ coordination and hydrogen bonding with Ala354, Glu384, and Tyr523 [10]. Quercetin acts by chelating the catalytic Zn^2+^ ion and forming hydrogen bonds with Glu384 and Tyr523, supported by its 3′,4′-dihydroxy B-ring and C2=C3 double bond [11]. Rutin, exhibits similar interactions with quercetin but with reduced affinity due to steric hindrance from its sugar moiety [12]. Hyperin modulates ACE by downregulating its expression in endothelial cells [13]. Among other metabolites identified, juncein (luteolin-4’-o-glucoside) —a glycosylated flavone— shares the core structure of luteolin known to inhibit ACE by mimicking classical inhibitors through chelation of Zn^2+^ via hydroxyl groupsm [11], it is likely that juncein retains ACE inhibitory activity.

Preliminary safety assessment was also conducted to determine appropriate dosing in subsequent pharmacological studies, PLAF was evaluated for acute oral toxicity at a limit dose of 2000 mg/kg body weight. No mortality or observable signs of toxicity were noted. Additionally, there were no significant changes in body weight, relative organ weights, gross necropsy findings, serum biochemical markers (SGPT, SGOT, BUN, creatinine, LDH), or histopathological alterations in vital organs (liver, heart, and kidney). These parameters indicate that the estimated *LD*_50_ of PLAF was greater than 2000 mg/kg based on the Globally Harmonized System (GHS), which is generally considered to have low acute toxicity; however, further sub-acute, chronic, and pharmacokinetic studies are recommended to establish its complete safety profile.

## CONCLUSION

This study demonstrated that *Psychotria luzoniensis* exhibits significant *in vitro* ACE I inhibitory activity, with PLAF and its subfraction PLAF-1D being the most potent inhibitors. Metabolite profiling of the bioactive fractions revealed the presence of flavonoids, including kaempferol, quercetin, rutin, and hyperin, all of which are known to elicit ACE inhibitory effects. In addition, several compounds including 4-(3,4-dihydroxyphenyl)-7-hydroxy-2-oxo-2H-chromen-5-yl-β-D-glucopyranoside, quercetin-3-O-β-D-xylopyranosyl(1→6)-β-D-glucopyranoside, cyanidin 3-O-lathyroside cation, juncein, loliolide, and butin were also detected which rarely reported in *Psychotria* species and are of particular interest for future phytochemical isolation and mechanistic studies. Acute oral toxicity testing at a limit single dose of 2000 mg/kg body weight revealed no adverse effects, supporting the safety of the extract for further biological evaluation. These findings suggest for further complete safety evaluation and *in vivo* assessment of the blood pressure-lowering activity of the active fractions to validate their safety and antihypertensive activity, respectively. Furthermore, exploration of additional antihypertensive mechanisms—such as modulation of the endothelial or sympathetic pathways —is recommended to provide a more comprehensive pharmacodynamic profile.

## ACKNOWLEDGMENTS

The authors gratefully acknowledge the support provided by the laboratories and research facilities that contributed to this study, particularly the Tuklas Lunas Development Center of Mariano Marcos State University, City of Batac, Philippines, and the Department of Pharmacognosy at Tohoku Medical and Pharmaceutical University, Sendai, Japan. Acknowledgement is also extended to Mr. Michael Calaramo, taxonomist at the Northwestern University Ecological Park in Laoag City, Philippines, for his valuable assistance in the collection and identification of plant samples.

## AUTHOR CONTRIBUTIONS

All authors significantly contributed to the study’s conceptualization, design, data acquisition, and/or data analysis and interpretation. Each participated in drafting the manuscript or critically revising it for substantial intellectual content, approved the final version for publication, and consented to submission to the present journal. Furthermore, all authors accept full responsibility for the integrity of the work and affirm their authorship eligibility in accordance with the guidelines of the International Committee of Medical Journal Editors (ICMJE).

## ETHICAL APPROVALS

All animal experiments were reviewed and approved by the Animal Experimental Committee of Tohoku Medical and Pharmaceutical University **(Approval No. A23073)**. The study was conducted in compliance with the university’s institutional ethical standards and international guidelines for the care and use of laboratory animals. Euthanasia of mice was carried out in accordance with the American Veterinary Medical Association (AVMA) Guidelines for the Euthanasia of Animals.

## DATA AVAILABILITY

All data generated and analyzed are included in this research article.

## PUBLISHER’S NOTE

All claims expressed in this article are solely those of the authors and do not necessarily represent those of the publisher, the editors, and the reviewers. This journal maintains neutrality regarding jurisdictional claims in published institutional affiliations.

